# A Pleiotropy-Informed Bayesian False Discovery Rate adapted to a Shared Control Design Finds New Disease Associations From GWAS Summary Statistics

**DOI:** 10.1101/014886

**Authors:** James Liley, Chris Wallace

## Abstract

Genome-wide association studies (GWAS) have been successful in identifying single nucleotide polymorphisms (SNPs) associated with many traits and diseases. However, at existing sample sizes, these variants explain only part of the estimated heritability [1]. Leverage of GWAS results from related phenotypes may improve detection without the need for larger datasets [2].

The Bayesian conditional false discovery rate (cFDR) [3] constitutes an upper bound on the expected false discovery rate (FDR) across a set of SNPs whose p values for two diseases are both less than two disease-specific thresholds. Calculation of the cFDR requires only summary statistics and has several advantages over traditional GWAS analysis. However, existing methods require distinct control samples between studies. Here, we extend the technique to allow for some or all controls to be shared, increasing applicability. Several different SNP sets can be defined with the same cFDR value, and we show that the expected FDR across the union of these sets may exceed expected FDR in any single set. We describe a procedure to establish an upper bound for the expected FDR among the union of such sets of SNPs.

We apply our technique to pairwise analysis of p values from ten autoimmune diseases with variable sharing of controls, enabling discovery of 59 SNP-disease associations which do not reach GWAS significance after genomic control in individual datasets. Most of the SNPs we highlight have previously been confirmed using replication studies or larger GWAS, a useful validation of our technique; we report eight SNP-disease associations across five diseases not previously declared.

Our technique extends and strengthens the previous algorithm, and establishes robust limits on the expected FDR. This approach can improve SNP detection in GWAS, and give insight into shared aetiology between phenotypically related conditions.

**Author Summary:** Many diseases have a significant hereditary component, only part of which has been explained by analysis of genome-wide association studies (GWAS). Shared aetiology, treatment protocols, and overlapping results from existing GWAS suggest similarities in genetic susceptibility between related diseases, which may be exploited to detect more disease-associated SNPs without the need for further data.

We extend an existing method for detecting SNPs associated with a given disease by conditioning on association with another disease. Our extension allows GWAS for the two conditions to share control samples, enabling larger overall control groups and application to the common case when GWAS for related diseases pool control samples. We demonstrate that our technique limits the expected overall false discovery rate at a threshold dependent on the two diseases.

We apply our technique to genotype data from ten immune mediated diseases. Overall pleiotropy between phenotypes is demonstrated graphically. We are able to declare several SNPs significant at a genome-wide level whilst controlling at a lower false-discovery rate than would be possible using a conventional approach, identifying eight previously unknown disease associations.

This technique can improve SNP detection in GWAS by re-analysing existing data, and gives insight into the shared genetic bases of autoimmune diseases.

## Introduction

Genome-wide association studies (GWAS) have enabled identification of genetic variants associated with a wide range of complex phenotypes, but in many cases these variants explain only a proportion of the known heritability [4]. There is increasing evidence that this is due to the combined contribution of small effects arising from multiple distinct variants [5]. The testing of a large number of potential variants in parallel, with a comparatively low number of samples, mandates a stringent threshold for significance in order to limit false positives (type 1 errors), meaning that discovery of variants responsible for small effects requires very large sample sizes. Detection of such variants by increasing numbers of samples in studies is time-consuming and expensive, particularly for rare phenotypes, but it may be possible to improve detection by re-analysis of existing data [6]. One promising strategy is to co-analyse GWAS results from similar phenotypes to exploit potential similarities in genetic aetiology. This has been attempted using several different methods [2, 7, 8].

The assumption that GWAS for similar diseases may yield overlapping sets of disease-associated variants is based on the phenomenon of pleiotropy, in which a genetic variant is associated with more than one trait or disease [9]. Pleiotropy is common in human genes: even when exclusively considering single nucleotide polymorphisms (SNPs) with strong evidence of association, around 15% of those associated with at least one trait are associated with multiple traits [10]. Elements of shared genetic aetiology may be suspected in diseases with similar symptomatology, such as bipolar disorder and schizophrenia [11] or in diseases with common risk factors, such as type 2 diabetes and obesity [12]. If two diseases are known or suspected to share associated genetic variants, a degree of association of a locus with one disease may increase the likelihood of association with the other. Use of external covariates in this way can alleviate some of the effect of multiple testing [8], meaning that phenotypic similarity may lead to improved detection of disease-associated variants. Correspondingly, discovery and specification of shared genetic aetiology between two diseases may suggest some shared pathophysiology [12].

A technique for improved discovery of disease variants using pleiotropy between pairs of diseases has been successfully developed and applied by Andreasson et al [3,13,14]. The technique extends the empirical Bayesian false discovery rate [15] to a two-phenotype scenario, in which association with one phenotype is tested conditional on varying degrees of association with another. We denote the phenotype for which association is being tested as the ‘principal phenotype’ and the other as the ‘conditional phenotype’.

By successively restricting attention to SNPs with a given strength of association in the conditional phenotype, the number of parallel tests to perform for association with the principal phenotype is reduced. If the two phenotypes share common associated variants, this restriction will retain disease-associated SNPs at a higher rate than null SNPs, resulting in a higher proportion of disease-associated SNPs in the restricted group than in the whole. The ‘conditional false discovery rate’ (cFDR), defined as the probability that a SNP is not associated the principal phenotype given its p values for the principal and conditional phenotypes are below some thresholds, exploits this effect. By computing cFDRs for schizophrenia conditioned on bipolar disorder and vice versa, Andreasson et al [3] identified multiple previously undiscovered loci for both. In a separate study computing cFDRs for hypertension conditioned on 12 related traits [13], 42 new loci associated with hypertension were reported. These constituted considerable improvement on existing results using single GWAS, albeit using a rather relaxed threshold of estimated *cFDR* ≤ 0.01.

A major disadvantage of the algorithm developed and used by Andreasson et al is the requirement that control groups for the two GWAS be distinct, in order to ensure that observed effect sizes are uncorrelated at null SNPs. This requires splitting a pool of potential controls between studies, with the summary statistics for each GWAS computed from only the controls allocated to that study. This may be impractical as it requires access to raw genotype data. More importantly, accuracy of effect size estimates improves with larger control groups, and consequently splitting controls in this way weakens the effect size estimates for individual studies. For this reason, many researchers employ a study design in which controls are pooled into a large group; for example, the Wellcome Trust Case Control and ImmunoChip consortia [16, 17].

Here we extend the cFDR approach to studies with overlapping control groups, exploiting an approach developed by Zaykin et al, following Lin et al [18,19] to adjust for the effect of shared controls. This allows the strongest available estimates of effect sizes to be used for calculation, and consequently strengthens the power of the technique. Our technique additionally allows cFDR rates to be computed from summary statistics alone, without the need to recalculate effect sizes after re-allocating controls. We demonstrate the improvement arising from sharing controls in a type 1 diabetes data set.

We also identify a previously undiscussed difficulty with the technique potentially leading to a falsely low estimate of false discovery rate amongst SNPs declared non-null. Multiple overlapping sets of SNPs may be defined each of which has *cFDR* ≤ *α*. However, the union of these sets does not necessarily have an expected false-discovery rate less than *α* and is generally higher. An implication of this is that if we declare non-null all SNPs for which estimated cFDR is less than *α*, the expected overall false-discovery rate amongst SNPs declared non-null is greater than *α*. We describe an upper bound on the false discovery rate amongst such SNPs based on areas of regions of the unit square.

We apply our method to summary SNP association statistics for ten phenotypically distinct autoimmune diseases: type 1 diabetes (T1D) [20], autoimmune thyroid disease (ATD) [21], coeliac disease (CEL) [22], multiple sclerosis (MS) [23], narcolepsy (NAR) [24], primary biliary cirrhosis (PBC) [25], psoriasis (PS) [26], rheumatoid arthritis (RA) [27], ulcerative colitis [28], and Crohn’s disease [28]. All were genotyped using a common SNP array: the ImmunoChip, designed to provide dense genotype coverage of regions associated with autoimmune disease. Many autoimmune traits are known to have significant heritability, much of which remains unexplained [29]. We hypothesised that our method can improve detection of disease-associated variants in these diseases without the need for distinct control groups.

## Results

### Overview of method

The unconditional false discovery rate for a set of SNPs with p values < *p_i_* is defined as the probability that a random SNP from this set is null. We denote this as *uFDR*(*p_i_*), and our estimate as 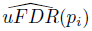.

The conditional false discovery rate (*cFDR*) is defined [3, 14] as the probability that a random SNP is null for a phenotype *i* given that the observed p values at that SNP for phenotypes *i* and *j* are less than (*p_i_*, *p_j_*); that is, 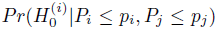, where 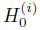 is the null hypothesis that the SNP is not associated with phenotype *i*. We denote this quantity as *cFDR*(*p_i_*|*p_j_*), and call phenotype *i* the ‘principal phenotype’ and phenotype *j* the ‘conditional phenotype’.

We first apply genomic control to allow the assumption that, globally, P values for null SNPs are uniformly distributed on [0, 1]. We compute an estimate of the *cFDR*, which we denote 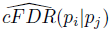, in a similar manner to that proposed by Andreasson et al, but incorporating expected non-uniformity in the distribution of *P_i_* due to the sharing of controls. As 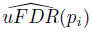 is monotonically related to *p_i_*, we set a significance cutoff at the maximum value of 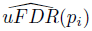 with *p_i_* < 5 × 10^−8^. Correspondingly, we set a significance cutoff for 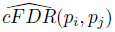 at the maximum 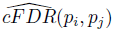 with *p_i_* < 5 × 10^−8^. Implementation of these steps in *R* is available from https://github.com/jamesliley/cFDR-common-controls.

### Sharing of control subjects

If no controls are shared between studies, it is reasonable to assume that observed effect sizes for the two phenotypes are independent under a null hypothesis for the principal phenotype. This implies that the expected quantile of a given SNP’s p value for the principal phenotype is simply the p value itself regardless of its p value for the conditional phenotype. However, when control samples are shared, this assumption is invalid. Shared controls induce a positive correlation on estimated effect sizes for the principal and conditional phenotype [18, 19], meaning that when attention is restricted to SNPs with a given degree of association with the conditional phenotype, the p values for the principal phenotype will be falsely low; that is, the probability 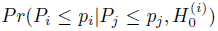 will not in general be equal to 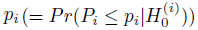; in fact it will usually be higher.

When controls are shared, the distribution of p values for the principal phenotype given p values for the conditional phenotype depends on the underlying effect of each SNP on the conditional phenotype. For any given SNP, this underlying effect size, which we denote *η*, is not known. However, across all SNPs, *η* may be considered to be realisations of a random variable *H* whose distribution is mirrored by the distribution of observed effect sizes for the conditional phenotype. By integrating over this unknown true effect size for the conditional phenotype, allowance can be made for shared controls, and the ‘expected quantile’ of a p value for the principal phenotype, defined as 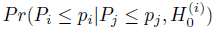, can be calculated, as detailed in the Methods section.

We assume that *H* has a mixture distribution defined by two parameters (*π*_0_, *σ*^2^), such that *H* = 0 with probability *π*_0_ and *H* ∼ *N*(0, *σ*_2_) with probability 1 − *π*_0_. The parameters (*π*_0_, *σ*) are estimated from the observed distribution of effect sizes for the conditional phenotype. In order to show the effect of our p value adjustment, we simulated p values for 20,000 SNPs for a principal and conditional phenotype, with controls shared between simulated studies. All SNPs were null for the principal phenotype, and were variably null or non-null for the conditional phenotype with probability 0.9, 0.1 respectively. *Z* scores at non-null SNPs for the conditional phenotype were distributed as *N*(0, *σ*^2^), as per our assumption. A value of *σ* = 3 was used, which was similar to the values of *σ* in real data estimated by our E-M algorithm.

We considered the set of simulated SNPs with p values for the conditional phenotype less than 0.05 (Figure 1). In the absence of shared controls, we expect the distribution of *p_i_* amongst this set to be uniform, and hence expect the black dots to lie along the x-y line. However, we see the principal p values are biased downward in this set (black dots, Figure 1). Our computed expected quantile (blue dots) agrees closely with the observed quantile. In a sense, this constitutes ‘adjusting’ the p values for the principal phenotype so that the expected distribution is uniform under the null hypothesis. Software to generate this simulation is available at https://github.com/jamesliley/cFDR-common-controls.

**Figure 1.**
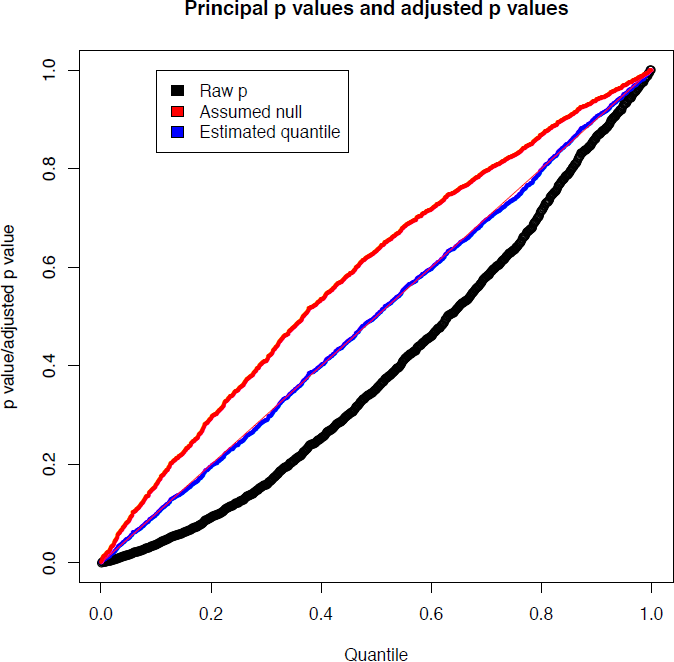
Correction for shared controls. Simulation of GWAS summary statistics for 20000 SNPs, all null for phenotype *i* and variably null or non-null for phenotype *j*, with association tested using a shared control group. Black dots show p values for phenotype *i* at all SNPs with p value for phenotype *j* less than 0.05, with evident downward bias. Blue dots show our adjustment to expected quantile of p values. The red dots show the expected quantile we would compute if we were to assume incorrectly that all SNPs were null for the conditional phenotype *i*. We see that this quantity overestimates the true quantile.

Our formula can easily be adapted to arbitrary distributions of *H* at the cost of increased computational time, but the form of the distribution of *H* is not generally known. We show in Text S1 (section A) that, even for distributions of *H* which differ markedly from our assumption of normality, the error in the estimate is not large, and generally translates to a negligible difference in the set of SNPs declared non-null using the 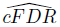 method. While our assumption has the potential to be anti-conservative if *H* is bimodal, nonparametric estimates for distributions of effect sizes suggest they have a uni-modal distribution centred on zero [30]. Reassuringly, our assumption is conservative if H has heavier tails than a normal.

### Comparison to split control approach

We compared Andreasson’s approach to SNP discovery which advocated splitting controls into non-overlapping subsets to our extended shared-control approach using a type 1 diabetes dataset with a total 12,175 cases and 15,171 controls. Controls and cases were each split into two sets (control sets had size 7,585 and 7,586, cases 6087 and 6088). ‘Split’ p values were computed using one set of controls and one set of cases and corresponding ‘shared’ p values were computed using the complete set of controls. As expected, more shared p values reached genome-wide significance than did split p values (Figure 2).

**Figure 2.**
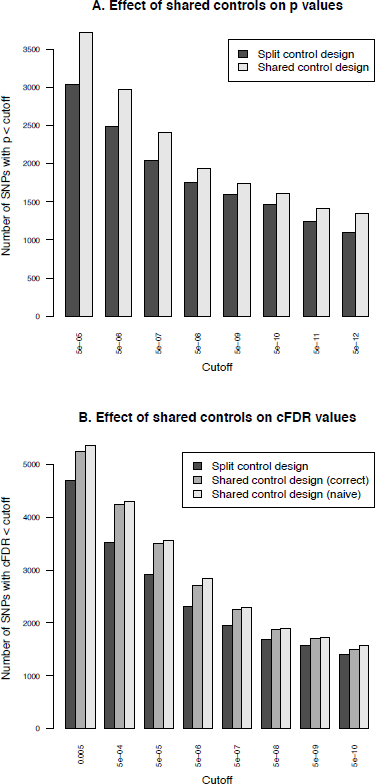
Validation of the shared-control approach and the p-value adjustment due to shared controls. Panel A shows the effect of splitting controls on power to detect association. The number of SNPs with p values less than a given cutoff are shown for split-control and shared-control approaches. For all p-value cutoffs, fewer SNPs reach significance when using a split-control design. Panel B shows the number of SNPs with 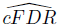 values less than a given cutoff using the existing method on a split-control design, our extended method on a shared control design with the adjustment for shared controls, or using the split-control approach naively on the shared-control design; that is, assuming incorrectly that 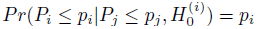. The second figure shows that failing to correctly calculate 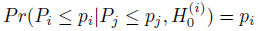 leads to a subtle increase in the number of SNPs declared non-null at all cutoffs, due to the incorrect underestimation of 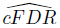.

We computed *cFDR* values by labelling one set of cases ‘conditional’ and the other ‘principal’ using the split-control p values using Andreasson’s approach and using the shared control p values using our method. For reference, we compared these to a naive application of Andreasson’s method on the shared-control p values (Figure 2B). More SNPs can be declared significant according to *cFDR* using the shared-control than split-control approach at all reasonable thresholds, and naive application of Andreasson’s approach to shared-control p values again increases the number declared significant.

Because the quantity 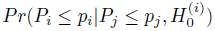 is systematically underestimated when using this naive method (by assuming it is equal to *p_i_*) as shown in Text S1 (section C), it leads to a falsely low 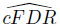. The increase in observed number of SNPs declared significant when using the naive method shows that it can indeed lead to false discoveries.

For principal phenotype p values in the range 5 × 10^−6^ - 5 × 10^−8^ - effectively the region from which ‘new’ SNPs may be discovered by cFDR rather than p value alone - the naive *cFDR* is frequently underestimated by 2-3 fold (Supplementary Figure S1, left panel). For lower p values, the naive *cFDR* may underestimate by hundreds- or thousand-fold, with the potential fold underestimation increasing with decreasing p value (Supplementary Figure S1, right panel). Because of the relatively high ratio of number of controls to number of cases, the correlation between effect sizes is lower in this constructed case (c. 0.22) than between most phenotypes in our study (c. 0.5). The underestimation of *cFDR* using the ‘naive’ method worsens with higher correlation, so we would expect that the fold-underestimate we see here is less severe than that which would be observed if applying this to other studies.

### An upper bound on the false discovery rate of all declared SNPs

An important property of our method is the control of the expected false discovery rate (FDR): the expected proportion of false positives among the SNPs found by our method. The p values at a SNP for the principal and conditional phenotype correspond to a point in the unit square. In this sense, we can define the expected FDR of a region *R* of the unit square as the ratio of the expected number of null SNPs whose p values are in *R* divided by the expected total number of SNPs whose p values are in *R*. From a result of Benjamini and Hochberg [31], the expected FDR when *R* is a rectangle with vertices at (0, 0), (*p_i_*, 0), (*p_i_*, *p_j_*), (0, *p_j_*) is at most 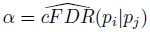. If we denote by *L* the closed region defined by the set of p value pairs *p_i_*, *p_j_* such that 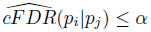, then *L* has the property that the FDR of any rectangle of this form contained within *L* is less than *α*.

However, the expected FDR over *L* is not necessarily bounded by *α* (Figure 3). This can be seen most easily in the extreme scenario in which all non-null SNPs are concentrated in the lower left corner of the unit square, and all null SNPs are also null for the conditional phenotype. In this case, the expected number of null SNPs in a rectangle is proportional to its area, so L is the union of all rectangles of the form above of a given area containing the lower-left corner of the unit square; that is, a hyperbola. Clearly the area of L is larger than the area of these constituent rectangles, yet it contains the same number of non-null SNPs, so it has a lower FDR.

**Figure 3.**
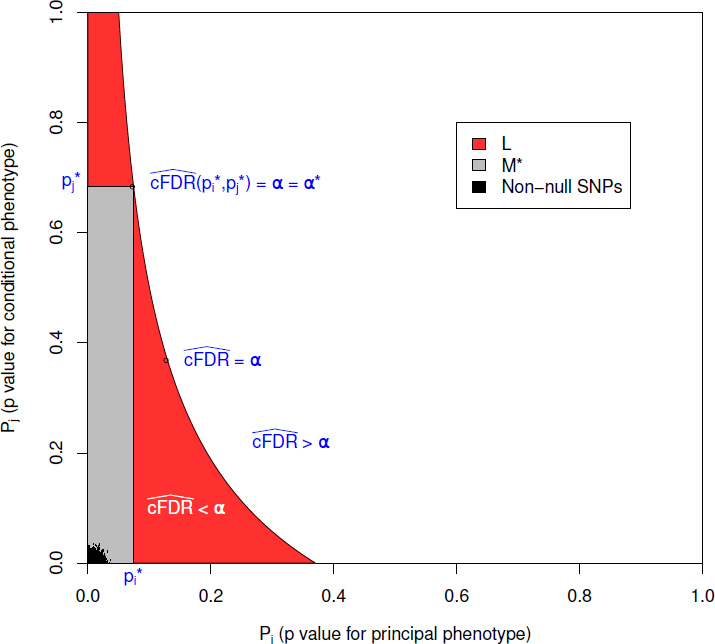
L is the locus of a set of points with 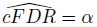. The FDR is the ratio of null SNPs to total SNPs in *L*. If all the non-null SNPs were concentrated in the lower left corner, then the number of non-null SNPs in *L* would be equal to that in any individual rectangle with vertices at the origin and on *L*, but the number of null SNPs would be greater, meaning that the expected FDR of all SNPs in *L* would be greater than *α*. *M*\* is the largest rectangle by area contained within *L*. The false discovery rate within *M*\* is less than *α*\*, the value of 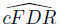 at the upper right vertex, which is usually equal to *α*, as in this case. The FDR of *L* is bounded by *α*\* υ(*L*)/ υ(*M*), where υ(*L*) and (*M*\*) are the expected number of null SNPs in *L* and *M*\* respectively (Text S1, section B).

In the original method [3], SNPs were declared significant if they were contained within any rectangular regions with a 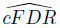 value of less than 0.01. Our reasoning demonstrates that the expected false-discovery rate amongst all such SNPs was higher than 0.01. We can derive an upper bound for the expected FDR of L by considering *M*\*, the largest rectangle in *L*. We show in Text S1 (section B) that the bound may be expressed simply as 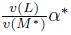, where *υ* () denotes the expected number of null SNPs contained within *L* or *M*\* (approximately the area of *L* and *M*\*) and *α*\* is the cFDR at the upper-right vertex of *M*\* (Figure 3).

### Application to ten immune mediated diseases

We obtained summary statistics in the form of p values for ten immune mediated diseases from ImmunoBase (www.immunobase.org, accessed 19/3/14). For each pair of diseases, the number of shared controls was estimated according to the description of the control samples in each paper. The numbers of cases, controls and our estimated numbers of shared controls for each study are shown in Table 1. Uniform quality control criteria were applied to all SNPs, and the MHC region, which exhibits both strong LD and strong association with immune mediated diseases was excluded. P values were corrected within each trait for genomic inflation using a standard algorithm [32] applied to SNPs included on the ImmunoChip to replicate a GWAS study of reading and maths ability (Steve Eyre and Cathryn Lewis, personal communication), unlikely to be related to any immune mediated disease studied here.

**Table 1.** Study sizes.

| Disease |  | Controls | Cases | Estimated number of pairwise shared controls |  |  |  |  |  |  |  |  |  |
| --- | --- | --- | --- | --- | --- | --- | --- | --- | --- | --- | --- | --- | --- |
|  |  |  |  | T1D | ATD | CEL | MS | NAR | PBC | PSO | RA | UC | CRO |
| T1D | [20]* | 12175 | 15171 | - | 9364 | 12228 | 8430 | 4289 | 8514 | 4822 | 8430 | 4020 | 4020 |
| ATD | [21] | 9364 | 2733 |  | - | 9364 | 8430 | 4289 | 8514 | 4822 | 8430 | 4020 | 4020 |
| CEL | [22] | 12228 | 12041 |  |  | - | 8430 | 4289 | 8514 | 4822 | 8430 | 4020 | 4020 |
| MS | [23] | 24091 | 14498 |  |  |  | - | 4289 | 8430 | 4822 | 8430 | 10102 | 10102 |
| NAR | [24] | 10421 | 1886 |  |  |  |  | - | 4289 | 4289 | 4289 | 4020 | 4020 |
| PBC | [25] | 8514 | 2861 |  |  |  |  |  | - | 4822 | 8430 | 4020 | 4020 |
| PSO | [26] | 22806 | 10588 |  |  |  |  |  |  | - | 4822 | 4020 | 4020 |
| RA | [27] | 15870 | 11475 |  |  |  |  |  |  |  | - | 4020 | 4020 |
| UC | [28] | 15977 | 10920 |  |  |  |  |  |  |  |  | - | 15977 |
| CRO | [28] | 15977 | 14763 |  |  |  |  |  |  |  |  |  | - |

Number of cases and controls for each study, and relevant references, together with our estimates of the number of controls shared between studies. \*P values for T1D are from a meta-analysis of case-control and TDT data, with effective numbers of cases computed as shown in the methods section.

P values for each principal phenotype were adjusted to *p*′ as described above in order to account for the effect of shared controls. For each ordered pair of phenotypes, a Q-Q plot was generated as per Andreasson et al [3]. A Q-Q plot is a graph of the observed distribution of a random variable against the expected distribution. We overlaid Q-Q plots for *log*_10_(*p*′) values for the principal phenotype for subsets of SNPs exhibiting successively smaller p values for the conditional phenotype. Figure 4 shows QQ plots for T1D conditional on RA and PSO; plots for all other pairwise comparisons may be found in Supplementary Figures S4-S13. Notably, if lines shift further left with more stringent cutoffs on association with the conditional phenotype, then SNPs which are associated with the conditional phenotype are more likely to be associated with the principal phenotype, indicating pleiotropic effects of SNPs on the two phenotypes. In many cases, the Q-Q plots demonstrate considerable leftward shift with conditioning on association with a second disease, and we see strong evidence for pleiotropy for T1D conditioned on RA and little or no evidence for pleiotropy for T1D conditioned on PSO.

**Figure 4.**
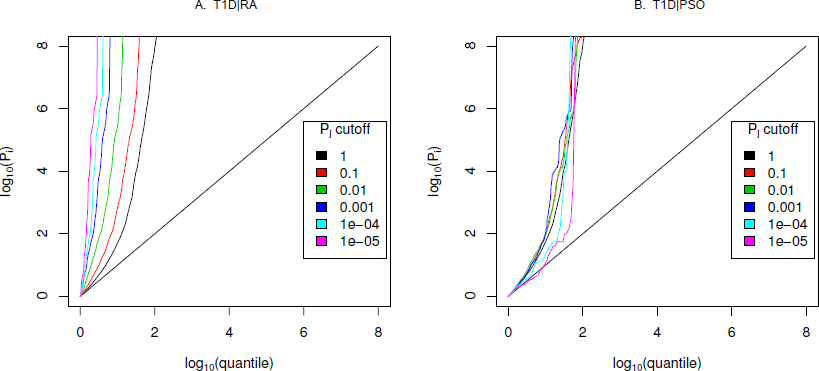
Q-Q plots for T1D conditional on RA (Panel A) and PSO (Panel B) Y axes show 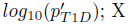; X axes show log quantile (rank) of p values in various sets of SNPs. The degree of leftward shift of a black point from the diagonal is proportional to the unconditional FDR of that p value for the principal phenotype, and the degree of leftward shift of a coloured point is proportional to the conditional FDR of the p value for the principal phenotype and the p-cutoff corresponding to the colour for the conditional phenotype. As expected, a leftward shift is seen even for the unconditional Q-Q plots (black line) owing to the use of the ImmunoChip, which focuses on potential autoimmune-associated regions. Each colour corresponds to the Q-Q plot for *p*_*T* 1*D*_ amongst a subset of SNPs with *p_RA_* or *p_PSO_* less than the indicated cutoff. P values for T1D are adjusted for the effect of shared controls between studies. A leftward shift with decreasing *p_RA_* or *p*_*PSO*_ cutoff indicates that SNPs which are associated with the conditional phenotype (RA or PSO) are more likely to be associated with the principal phenotype (T1D), presumably due to pleiotropic effects on phenotypes. Good enrichment is seen for T1D conditioning on RA (Panel A), and little or no enrichment conditioning on PSO (Panel B).

We estimated the unconditional and conditional false discovery rates, 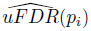 and 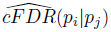, at each SNP for each phenotype and each ordered pair of phenotypes respectively. Figure 5 shows 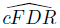 for T1D conditioned on RA. The advantage gained by 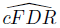 can be seen in the left-shift of the region in which a SNP can be declared significant (blue dots), corresponding to a higher p-value cutoff for significance for T1D among SNPs with low p values for RA. Indeed, if only SNPs with a p value for RA less than some threshold ζ are considered, a p value cutoff for significance for T1D is given by the leftmost border of the blue dots on the line *P_j_* = ζ.

**Figure 5.**
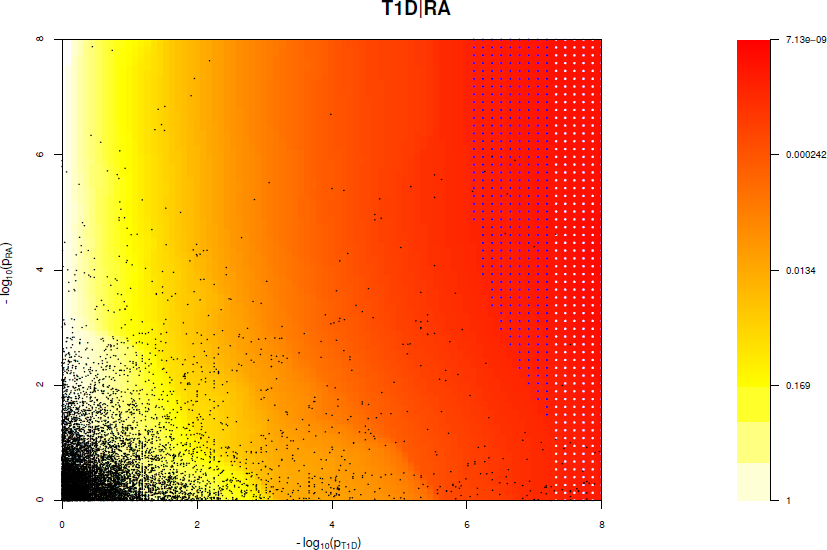
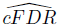 (red-yellow) for T1D conditioned on RA. White dots signify the region for which 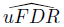 is less than *α* corresponding to *p* < 5 × 10^−8^. Blue dots signify the region for which cFDR is less than the same *α*. Note the leftward shift of blue points and the general leftward shift of colours corresponding to an increased p-value threshold for association with T1D for SNPs with low p values for RA. Black dots show a random sample of the observed p value pairs.

The degree of leftward shift in the Q-Q plots clearly contains information about the degree of pleiotropy between diseases. We defined a statistic summarizing some aspects of this evidence for pleiotropy and used it to visualise the set of pairwise relationships between diseases as a network (Figure 6). The network encouragingly reflects several pathophysiological associations: UC is linked to CRO, and T1D to ATD. Strong linkage is also seen both ways for MS and PBC, and between T1D and RA, findings which can also be seen in the Q-Q plots (Supplementary Figures S4–S13). One way relationships suggest the presence of a larger total number of associated SNPs for the disease at the start of the arrow than at the end.

**Figure 6.**
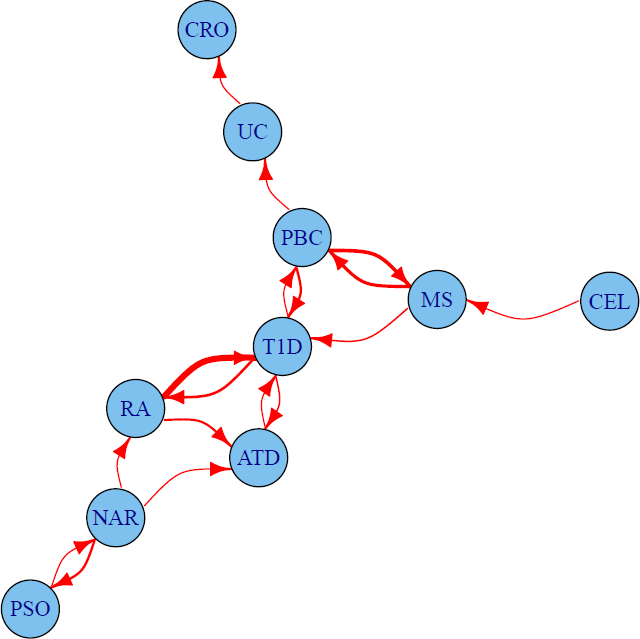
Network of degree of pleiotropy between phenotypes. An arrow runs from vertex *i* to vertex *j* if and only if by conditioning on *p* < 5 × 10^−6^ for the conditional phenotype *j* we can increase the threshold for significance for the p value for the principal phenotype *i* from 5 × 10^−8^ to 4 × 10^−7^ or greater. Edges are thickened if the cutoff could be increased more than this. The threshold 4 × 10^−7^ was selected as the minimum value for which the network is weakly connected; that is, having an arrow to or from each edge.

### Discovery of novel associations

The numbers of SNPs deemed significant for each phenotype by analysis using unconditional and conditional approaches are shown in table 2, with details in Supplementary Tables S2–S11. 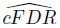 allows certain SNPs with p values as high as 3 × 10^−6^ to be declared significant while controlling the false discovery rate at a relatively low value. Fifty-one of the 59 SNPs we identify uniquely through 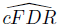 have previously been reported to be associated with the relevant disease through use of alternative significance thresholds, other genomic control procedures, other GWAS or additional samples not genotyped by ImmunoChip, a useful verification of our technique. Eight of the SNPs we discover uniquely through cFDR were in regions not previously known to be associated with the corresponding disease (table 3). These will require replication in independent samples to be declared truly associated, but they contain some potentially interesting signals, such as an association for RA at SNP rs72928038 near existing MS, ATD and T1D associations in *BACH2*, a transcriptional regulator involved in transcription repression and activation by *MAFK* [33]

**Table 2.** Number of association signals found by unconditional and conditional methods

|  | Univariate |  | Conditional |  | Efficiency ratio |
| --- | --- | --- | --- | --- | --- |
|  | N.SNPs | FDR | N.SNPs | FDR |  |
| T1D | 44 | $3.79 \times 10^{-6}$ | 4 | $7.32 \times 10^{-6}$ | 0.11 |
| ATD | 3 | $1.04 \times 10^{-5}$ | 4 | $1.02 \times 10^{-5}$ | 0.03 |
| CEL | 46 | $3.97 \times 10^{-6}$ | 7 | $5.77 \times 10^{-6}$ | 0.24 |
| MS | 43 | $5.97 \times 10^{-6}$ | 12 | $1.19 \times 10^{-5}$ | 0.15 |
| NAR | 3 | $1.61 \times 10^{-4}$ | 0 | $3.05 \times 10^{-4}$ | 0.83 |
| PBC | 25 | $8.86 \times 10^{-6}$ | 2 | $2.09 \times 10^{-5}$ | 0.12 |
| PSO | 44 | $2.33 \times 10^{-5}$ | 2 | $5.45 \times 10^{-5}$ | 0.06 |
| RA | 16 | $2.04 \times 10^{-5}$ | 10 | $6.00 \times 10^{-5}$ | 0.11 |
| UC | 49 | $4.49 \times 10^{-6}$ | 6 | $6.73 \times 10^{-6}$ | 0.14 |
| CRO | 80 | $2.08 \times 10^{-6}$ | 12 | $2.73 \times 10^{-6}$ | 0.28 |

Number of SNPs with $p \leq 5 \times 10^{-8}$ after genomic found by analysis of the principal phenotype alone and the estimated equivalent FDR for this set of SNPs. Conditional analysis shows the number of additional SNPs found through conditional FDR analysis, and the upper bound for the FDR of all SNPs selected by this method, including adjustment for the multiple phenotypes conditioned upon. Finally, we summarise the performance of the cFDR approach by the FDR ratio - the ratio of the upper bound for the FDR for all SNPs selected by the cFDR approach, and the estimated FDR of all SNPs with $p$ less than the maximum $p$ value within this set if we had not conditioned. Note, we list only the most associated SNPs in each LD block by pruning according to LD, as described in Methods.

**Table 3.** Novel SNP-disease associations.

| Chr | Pos | SNP | Disease | p value | C. phen. | MAF | Nearby Genes |
| --- | --- | --- | --- | --- | --- | --- | --- |
| 1p31.3 | 61791863 | rs6691768: A>G | CEL | $8.56 \times 10^{-8}$ | CRO | 0.373 | *NFIA* |
| 1p13.1 | 117076399 | rs1034920: T>C | PBC | $1.43 \times 10^{-6}$ | MS | 0.100 | *CD58* |
| 2p21 | 43359275 | rs6705577: G>C | CEL | $3.61 \times 10^{-7}$ | MS | 0.272 | *THADA* |
| 2q32.1 | 185501065 | rs79248157: T>C | UC | $1.53 \times 10^{-7}$ | CRO | 0.031 | *ZNF804A* |
| 6q15 | 90976768 | rs72928038: G>A | RA | $5.89 \times 10^{-7}$ | T1D | 0.176 | *BACH2* |
| 16q12.1 | 51080214 | rs12924003: C>T | CRO | $5.82 \times 10^{-8}$ | NAR | 0.321 | *NOD2* |
| 20q13.12 | 44596207 | rs6032606: G>C | CEL | $1.31 \times 10^{-7}$ | MS | 0.052 | *ZNF355, MMP9* |
| 22q13 | 37633851 | rs9610686: G>A | CEL | $1.90 \times 10^{-7}$ | ATD | 0.387 | *RAC2* |

Eight SNP associations with the indicated disease discovered by $cFDR$ but not previously published to our knowledge, together with the phenotype upon which they have been conditioned (C. phen.) and nearby genes. SNPs are shown by RSID: major>minor alleles. The disease (principal phenotype) p value has been corrected for genomic inflation. Note that a SNP reaching significance by $\widehat{cFDR}$ for the principal phenotype does not constitute evidence of association with the conditional phenotype (C. phen.). Chr = chromosome, Pos = position (build GRCh37), SNPs are shown with major<minor alleles, MAF = minor allele frequency in UK controls.

The SNP rs1034290 in region 1p13.1, which we found to be associated with PBC, is in intron three of *CD58*, which is a surface receptor involved in binding and activation of T-lymphocytes. The protective effect of the MS-associated allele is postulated to arise from upregulation of the transcription factor FOXP3 [34] and the patterns of association in the region suggest the two diseases may share a causal variant here (http://www.immunobase.org).

## Discussion

We have extended a technique for computing conditional Bayesian False Discovery Rates to GWAS for independent diseases with shared control groups. This technique enables improved detection of disease-associated SNPs compared to conventional methods. By enabling larger control groups for each study, our method uses data more efficiently than in corresponding study designs in which control groups are independent, and is applicable to a wider range of GWAS datasets for which only summary statistics are available.

Combination of GWAS by analysis of pleiotropy in this sense has several attractive advantages over single-phenotype analysis. The most obvious advantage is improved detection of disease-associated SNPs using GWAS without the need for additional samples. A secondary advantage arises from understanding of the pleiotropic structure between phenotypes: if a SNP is known to exhibit pleiotropy between two conditions, it may be causative for a shared risk factor or pre-disease state. Analysis of such SNPs has the potential to yield information on disease aetiology, with implications for preventative medicine and development of treatment.

A further potential use for this technique could be the genomic analysis of diseases with complex phenotypes. In many cases, distinction between two diseases may be difficult; for instance, Crohn’s disease and Ulcerative Colitis [35]. Additionally, many diseases, including narcolepsy (http://www.uptodate.com/contents/clinical-features-and-diagnosis-of-narcolepsy, accessed 20/6/14), are definitively diagnosed on clinical grounds. This implies that these diseases may constitute a range of biochemical and genetic states. Inclusion criteria based on objective biochemical grounds, such as that used for narcolepsy in the context of this paper [24] are unlikely to characterise all patients with these diseases, and conclusions drawn from studies will not necessarily be medically applicable to the whole patient population. Given this, diseases defined phenotypically with potential genomic diversity may be better analysed by separate consideration of biochemically-defined subtypes, with a collective analysis performed by a method such as 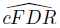, avoiding the assumption that the genomic bases of disease subtypes are identical.

We identify a counter intuitive property that the FDR in the union of all regions with 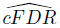 less than *α* given may be greater than *α*, and propose a method to overcome this problem. Our methods for adjusting cutoffs to control FDR and account for multiple testing demonstrate the geometrical elegance of the theory of these techniques, with the possibility for further improvements and understanding. They are complex to apply, but could be much simplified if interest was directed to SNPs with conditional p values less than some threshold *p_0_*. Our method would ensure that the expected false discovery rate at SNPs with 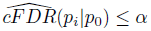 would indeed be controlled at *α*. Our more complicated method to control FDR is necessary if the variable *p_j_* is used in place of the constant *p_0_*.

An important consideration in both our method and the original Andreasson method is that a 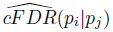 value which reaches significance does not constitute genome-wide evidence of association with the conditional phenotype *j*; indeed, the probability of association with the conditional phenotype relates to 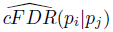 and in general 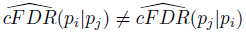. In some cases, where the principal p value is very close to genome-wide significance, even conditioning on *p_j_* ≤ 0.5 can theoretically be enough to reach the relevant 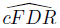 threshold. This is not a weakness of the 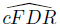 method as such, but a consequence of using a discrete technique (a significance cutoff) on a variable which essentially continuous in two dimensions 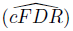 Principal p values greater than 5 × 10^−8^ which can be declared significant conditioning on large conditional p value cutoffs correspond to an increase in the area of the region *L* (see results section), which is accounted for by our FDR-controlling method.

Our method enables improved detection of SNPs compared to analysis of unconditional FDR (principal p value alone). However, the improvement is smaller than that reported by Andreasson et al [3,13,14], who detected almost twice as many SNPs using *cFDR* as they would have detected with *uFDR*. This is expected for two reasons. Firstly, the gain in power from *cFDR* essentially comes from an increase in the total number of controls and the effective number of cases. If controls are shared, the only information gain can come from increasing the number of effective cases. Consequently, the difference in power between *cFDR* and *uFDR* will not be as large when controls are shared, although both outperform their counterparts when controls are split. Secondly, we were careful to use stringent cutoffs for FDR which were chosen to mirror the established genomewide significance threshold of *p* ≤ 5 × 10^−8^, generally equivalent to a false discovery rate around 5 × 10^−6^ to 5 × 10^−5^, compared to Andreasson et al who declared non-null all SNPs with 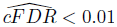.

One alternative way to exploit pleiotropic relationships is by meta-analysing two related diseases together, as though the diseases were the same. Our method confers several advantages over this approach. The most important of these is that our method borrows strength from other SNPs according to the level of genome wide pleiotropy between diseases; that is, if the two GWAS suggest extensive pleiotropy (such as Figure 4 for T1D — RA), a low p value for a conditional phenotype will ‘sway’ our judgement of association with the principal phenotype more than the same p value for a conditional phenotype with poor pleiotropy (such as Figure 4, for T1D — PSO). A meta-analysis would not distinguish these two scenarios. A secondary advantage of our technique is that SNP detection is not systematically weakened if the two diseases do not exhibit pleiotropy, as would be the case in meta-analysis; this arises because we are testing association with only one of the two phenotypes at a time.

## Methods

### Ethics Statement

This paper re-analyses previously published datasets. All patient data were handled in accordance with the policies and procedures of the participating organisations.

### Datasets

We obtained SNP summary statistics from ten studies on autoimmune diseases from ImmunoBase (www.immunobase.org). Inclusion and exclusion criteria for the studies are described in detail in the original publications ( [20–28,28]. Generally, some or all controls from different studies were obtained from common data sources, resulting in overlapping control groups. All studies used the ImmunoChip array [17].

P values for type 1 diabetes were from a meta-analysis of a case-control study and familial study using the transmission disequilibrium test (TDT). In order to calculate the correlation between p values for different diseases, we needed to calculate effective numbers of cases and controls for the combined T1D study. For a case control study, under the assumptions of Hardy-Weinberg and the null hypothesis, the variance of the log odds ratio may be expressed as

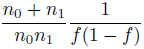

where *n*_0_ and *n*_1_ are the numbers of cases and controls and *f* is the minor allele frequency in controls.

Given the standard error of a log OR for the TDT study, and a minor allele frequency, we estimated *M* = *σ̂*^2^*f* (1 − *f*) for all ImmunoChip SNPs which did not show deviation from the null hypothesis (*p* > 0.5). The distribution of log(*M*) is shown in Supplementary Figure S3. By equating the median of *M* with 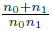, and assuming that each TDT family contributed the equivalent information to one control in a case-control study, ie *n*_0_ = 2943, we estimated an equivalent number of cases as 4126. This seemed reasonable, given that there are a total of 5505 (dependent) cases across those families.

SNPs were excluded on the basis of QC summaries calculated on 12,888 common controls: call rate less than 99%, minor allele frequency less than 0.02, or deviation from Hardy-Weinberg equilibrium (|*Z*| > 5). Given the strong association of immune mediated diseases with the MHC and the extended LD in the region, we were concerned that MHC SNPs might cause inaccurate estimation of pleiotropy. We therefore excluded SNPs in a wide band around the MHC region on chromosome 6 (co-ordinates 24500000: 34800000, build NCBI36). After quality control, genotype data was available for at least one phenotype at a total of 110677 SNPs.

### Genomic control

P values were corrected for genomic inflation using a genomic control algorithm [32]. A set of SNPs known to be unassociated with autoimmune disease was obtained from the Wellcome Trust Case Control Consortium (WTCCC) study on reading and mathematics ability. These SNPs were pruned so that none were in LD with *r*^2^ > 0.2, and any SNPs within 500 kb of known autoimmune-associated regions were removed. The average degree of inflation was computed for each disease at the remaining 1761 SNPs, and all effect sizes and p values were adjusted accordingly.

### Procedures for uFDR and cFDR

We assume that the p-values for a phenotype *i* across all SNPs are instances of a random variable *P_i_*. If *p_i_* is an instance of this random variable corresponding to a SNP of interest, the unconditional false discovery rate *uFDR*(*p_i_*) is defined as

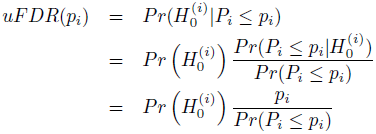

where 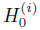 is the null hypothesis that the SNP of interest is not associated with phenotype *i*. Given a set of observed p values 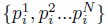 for a phenotype *i* at *N* different SNPs, and an observed p value *p_i_* for a SNP of interest, we estimate this quantity as

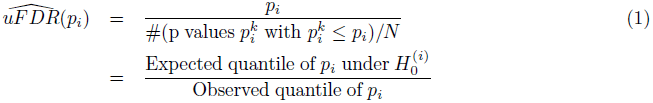

Because we make the approximation 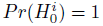, the estimate 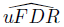 is a an upwards-biased estimate of *uFDR*; that is, its expected value is greater than the true *uFDR*, making it a conservative estimator.

We compute the quantity (1) for each SNP at each phenotype, declaring any SNP for which 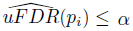 as non-null for phenotype *i*. Defining *V* as the number of SNPs falsely declared non-null, *R* as the total number of SNPs declared non-null, and *Q* = *V/R*, a theorem of Benjamini and Hochberg [31] shows the expected false discovery rate *E*(*Q*) among SNPs with 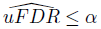 is less than *α*.

The *cFDR* constitutes a natural extension of this idea. We assume that the p-values for two phenotypes *i* and *j* across all SNPs are instances of a pair of random variables *P_i_*, *P_j_*. If *P_i_* and *p_j_* are instances of these variables corresponding to a SNP of interest then the conditional false discovery rate *cFDR* is defined for the set of SNPs with p values for each phenotype less than or equal to those at this SNP (as per Andreasson et al [3]) as

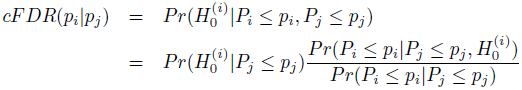

The estimation of this quantity proceeds in a similar way to *uFDR*. Given a set of observed p value pairs 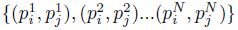 for two phenotypes *i* and *j* at *N* different SNPs, and an observed p value pair (*p_i_*, *p_j_*) for a SNP of interest, we define *N*_1_ as the number of p value pairs with *P_j_* ≤ *p_j_*, and estimate the *cFDR* as

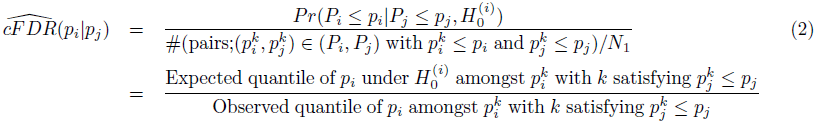

Again, this estimate is conservative, due to the approximation 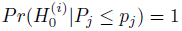.

We compute the quantity (2) for each SNP at each pair of phenotypes, declaring any SNP for which 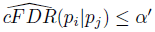 as non-null for phenotype *i*. However, as noted earlier, this does not guarantee that the expected false discovery rate amongst such SNPs is less than *α*′. We show that the FDR is controlled at a higher level dependent on the region of the unit square defined by rectangles for which 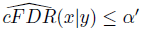.

Our method here diverges from the original method proposed by Andreasson et al, in the use of the expected quantile 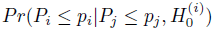 in place of the p-value *p_i_*. If studies share no controls, it can be reasonably assumed that, for a SNP which is null for phenotype *i*, the p values (*p_i_*, *p_j_*) are independent, so 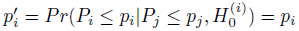. This is the approach taken by Andreasson et al [3]. We propose a method for computing 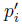 when controls are shared between studies, and the independence assumption above is not valid.

Our approach is to compute the related quantity 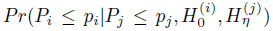, where *η* is the (unobserved) effect size we would observe for a given SNP for phenotype *j* if the observed MAFs agreed exactly with the population MAFs for that SNP, and 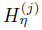 is the hypothesis that *Z_j_* ∼ *N*(*η*, 1) for that SNP. This quantity can be thought of as the ‘expected quantile’ of *p_i_*; that is, the proportion of p values we expect to be less than *p_i_*.

### Computation of expected quantile

From the first part of (2), we have:

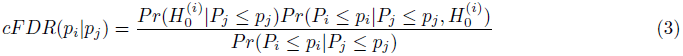

As per Andreasson et al [3], the quantity 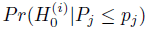 is set conservatively at 1, and the quantity *Pr*(*P_i_* ≤ *p_i_*|*P_j_* ≤ *p_j_*) is estimated empirically as the proportion of pairs of observed p values 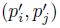 with 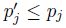 which also satisfy 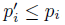.

For a given SNP, let *η* denote the standardised mean allele frequency (MAF) difference; that is, the *Z* value we would compute if the observed MAFs agreed exactly with the population MAFs. We consider *η* for a random SNP as being an instance of a random variable *H*, and that the observed *z* value for that SNP *Z|H* = *η* is distributed as

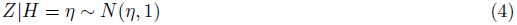

We further assume that *H* follows a mixture distribution taking the value 0 with probability 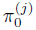 and a normal *pdf* with probability 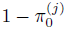

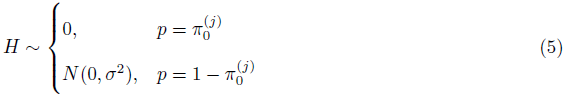

This implies

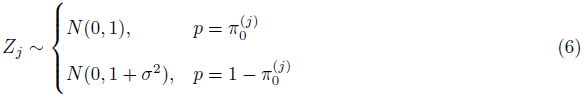

Thus, given the observed distribution of *Z_j_*, the parameters 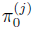 and *σ_j_* may be estimated by an expectation - maximisation algorithm (https://gist.github.com/chr1swallace/11421212).

We assume as per Zaykin [18] that the distribution of pairs of observed z values (*Z_i_*, *Z_j_*) for a single given SNP is bivariate normal. Denote by 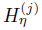 the event that, for a given SNP, the values *Z_j_* are distributed as *N*(*η*, 1), with n depending on the SNP.

Under our assumption of the null hypothesis 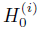 for the principal phenotype and a population MAF difference corresponding to *η* for the conditional phenotype, we have

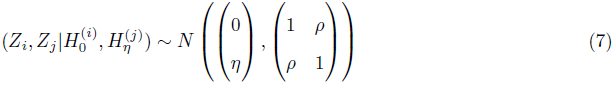

The correlation *ρ* arises from the shared controls between groups [18,19] and is asymptotically equal to

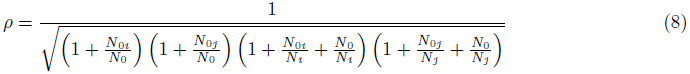

where *N_i_* and *N_j_* are the numbers of cases, *N*_0*i*_ and *N*_0*j*_ are the numbers of non-shared controls, and N_0_ is the number of shared controls for the original GWAS for the principal and conditional phenotypes respectively. There is good agreement with the asymptotic correlation when group sizes are greater than 100 [18].

Given equations (5)-(8), the joint distribution of *Z_i_* and *Z_j_* can be computed under only the assumption 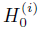. The value of the partial PDF of 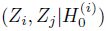 at (*x*, *y*) can be derived in a similar way to (6):

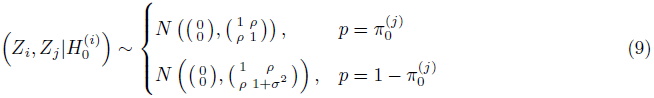

We now compute the final probability in equation (3). Define

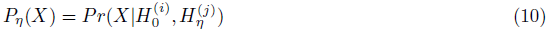

as the probability of observing events *X* for a particular SNP with true effect size *η* (which may be 0, corresponding to the general null). Then,

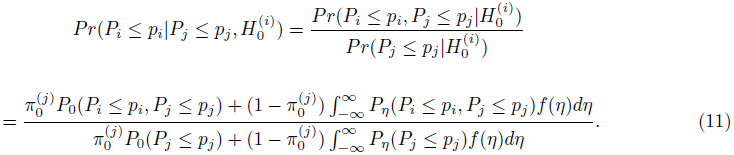

If the distribution of *H* is estimable by other means, quantity (11) can be calculated numerically without the assumption that the non-null component of *H* be normally distributed, at the cost of higher computation time. Under our assumptions, equations (6) and (9) enable the fast computation of quantity (11) by normal CDFs; writing

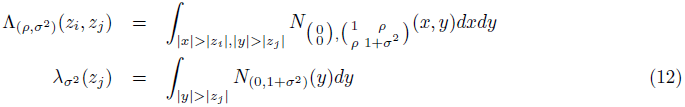

we have

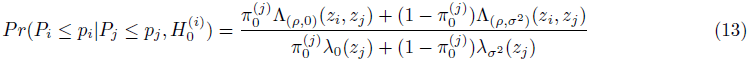

### Point expected quantile

Because the formula for 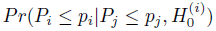 is differentiable on the unit square, an expression for the expected quantile of *p_i_*. given an exact value for *p_j_* can be computed by taking the partial derivative with respect to *p_j_*:

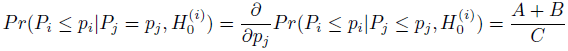

Where

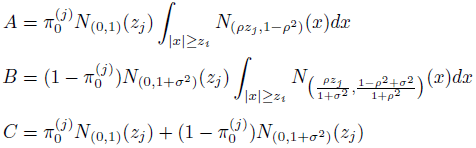

where *N*_*μ*,*σ*_^2^ (*x*) denotes the value of the normal *pdf* with mean and *μ* variance *σ*^2^ at *x*.

### Significance threshold

Because 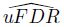 values are monotonically related to p values, the widely accepted GWAS p value cutoff of 5 × 10^−8^ corresponds naturally to a cutoff for 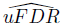. For each phenotype *i*, we set a significance threshold *β^i^* for 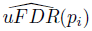 as the lowest possible value of γ for which 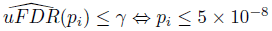.

We then applied an analagous approach to 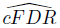 For each pair of phenotypes (*i, j*), we set a significance threshold 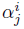 as the lowest possible value of γ′ for which 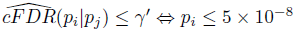. Given the distribution of *P_j_*, it is possible that this could lead to declaring SNPs with *p_i_* > 5 × 10^−8^, *p_j_* ≈ 1 as significant. To avoid this, if 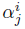 was larger (less stringent) than *β^i^*, we set 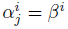.

For each ordered pair of phenotypes (*i,j*), we declared all SNPs with 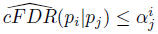 as non-null for phenotype *i*. This included all SNPs with 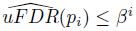. We then used a technique described in Text S1 (section B) to compute upper bounds 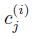 on the false discovery rate amongst SNPs for which 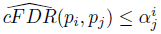. For each phenotype, this gave nine upper bounds, corresponding to each of the nine conditional phenotypes.

### Network and heatmap representation of pleiotropy

We compared the degree of pleiotropy between diseases by considering how much the p-value threshold for significance for the principal phenotype changed when conditioning on a small p-value threshold for the conditional phenotype. We used the 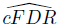 algorithm to compute the number 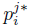 such that 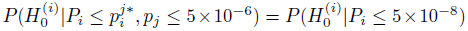; that is, 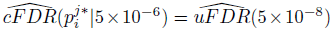. We then considered the ratio 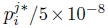; that is, the fold increase in significance cutoff after conditioning.

We note that because of the fixed value of *p_j_* =5 × 10^−6^, the expected false discovery rate amongst the set of SNPs which satisfy 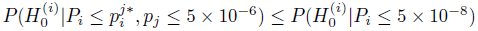 is bounded above by 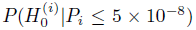, by the Benjamini-Hochberg result. Thus the expected false discovery rate amongst SNPs with 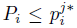 and *P_j_* ≤ 5 × 10^−6^ is bounded by the same value as the expected false discovery rate amongst SNPs with *P_i_* ≤ 5 × 10^−8^.

We visualised the ratio 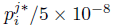 as a heatmap (Supplementary Figure S2). We also produced a network (Figure 6), with an edge from vertex *i* to vertex *j* if and only if, by conditioning on *P_j_* ≤ 5 × 10^−6^, the cutoff for significance for *P_i_* could be increased from 5 × 10^−8^ to 4 × 10^−7^. This cutoff was chosen as the smallest value such that the network was weakly connected; that is, each vertex had an arrow either to it or from it.

### Discovery of novel SNP associations

Cutoffs *β^i^* were chosen such that 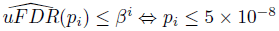. SNPs were deemed significant for a principal phenotype *i* if 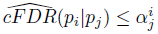 for any conditional phenotype *j* and 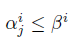.

The expected false discovery rate amongst SNPs for which 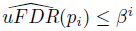 is less than *β^i^* due to a theorem of Benjamini and Hochberg. However, as discussed above and in Text S1 (section B), the expected false discovery rate amongst SNPs for which 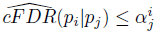 is not necessarily lower than 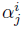. For each ordered pair of phenotypes (*i*, *j*), an upper bound 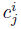 was computed for the expected false discovery rates amongst SNPs with 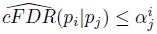. The list of SNPs declared non-null for phenotype *i* was pruned to allow for linkage disequilibrium (LD) by listing all SNPs in increasing order of 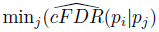 and stepping through the list from left to right, at each stage removing all SNPs in LD with *r*^2^ ≥ 0.1 to the right of the current SNP. This ideally leads to the inclusion of at most one SNP from each LD block.

### Multiple Testing

A multiple testing problem arises from considering p values for one disease conditioned separately on nine others. Specifically, if the criterion for declaring a SNP non-null for phenotype *i* is that 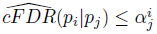 for at least one of the nine possible values of *j*, then the FDR for all SNPs declared non-null will be greater than the FDR among the smaller set of SNPs for which 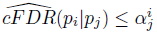 for only one value of *j*, due to multiple testing.

However, this excess FDR is not enough to warrant a Bonferroni (Sidak) correction; the 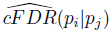 values for a phenotype *i* are highly correlated, as all are in turn highly correlated with *p_i_*. A Bonferroni correction tends to remove any advantage in SNP detection gained from 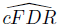, though an advantage may still be seen when only considering one conditional phenotype *j*.

We opted to use a method proposed by Nyholt [36] to correct for multiple testing in SNPs with high LD. We estimated a correlation matrix Ω for potentially non-null cFDR values using Spearman’s rank correlation. The variance of the eigenvalues of Ω, *Var*(λ_*obs*_), was computed and used to estimate the effective number of variables *M_eff_* according to the equation

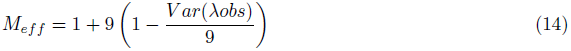

Note that *Var*(λ_*obs*_) is between 9 (completely correlated variables, effectively one test) and 0 (completely uncorrelated variables, essentially a Bonferroni correction).

Denoting by 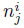 the number of SNPs with 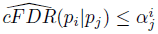, corresponding to an upper bound on the FDR of 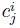, an upper bound for the FDR among all SNPs declared significant for phenotype *i* was then computed as

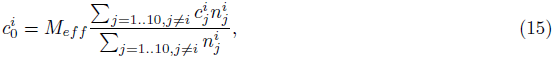

intuitively, multiplying the expected average number of false discoveries across conditional phenotypes 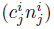 by the effective number of tests. Values of *M_eff_* and 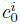 are shown in Supplementary Table S1.

## Acknowledgments

We thank Professor John Todd for help in interpreting the results. Thanks to Professor Cathryn Lewis and Dr Steve Eyre for the list of SNPs used for genomic control. We gratefully acknowledge the following groups and individuals who provided biological samples or data for this study. The IIBDGC shared summary association data on Crohn’s and ulcerative colitis. We would like to thank the UK Medical Research Council and Wellcome Trust for funding the collection of DNA for the British 1958 Birth Cohort (MRC grant G0000934, WT grant 068545/Z/02). We thank The Avon Longitudinal Study of Parents and Children laboratory in Bristol and the British 1958 Birth Cohort team, including S. Ring, R. Jones, M. Pembrey, W. McArdle, D. Strachan and P. Burton, for preparing and providing the control DNA samples. We acknowledge use of DNA from The UK Blood Services collection of Common Controls (UKBS collection). The collection was established as part of the Wellcome Trust Case-Control Consortium. We acknowledge use of DNA samples from the NIHR Cambridge BioResource. We thank volunteers for their support and participation in the Cambridge BioResource and members of the Cambridge BioResource SAB and Management Committee for their support of our study. Access to Cambridge BioResource volunteers and their data and samples is governed by the Cambridge BioResource SAB. Documents describing access arrangements and contact details are available at http://www.cambridgebioresource.org.uk/.

Acknowledgements for the individual studies whose summary data we accessed via ImmunoBase may be found at http://www.immunobase.org/poster/immunochip-paper-acknowledgments/.

## Supporting Information

**Text S1** Mathematical basis for estimation of cFDR and establishment of upper bound on expected FDR when using 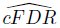 method.

**Table S1.** Calculation of false discovery rates for SNPs reaching 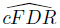 significance levels. *M_eff_* gives the ‘effective number of tests’, relating to the multiple testing adjustment for the multiple phenotypes conditioned upon (see Methods section). Max p is the maximum principal p value at which a SNP was able to be declared significant using 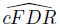. Eq FDR shows the false-discovery rate we would be forced to control at in order to detect all these SNPs the principal p value alone. The FDR bound (bold) is the false discovery rate at which the set of SNPs discovered by the 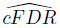 method is controlled.

**Table S2.** Tables S2–S11 show associated SNPs for each phenotype, ordered by best cFDR. P values shown are after adjustment for genomic inflation. Chromosome positions are from the NCBI36 assembly. The conditional phenotype shown is the phenotype for which the cFDR was most below the relevant cutoff. Column CP is the conditional phenotype for which corrected cFDR was lowest. SNPs with p value greater than 5 × 10^−8^ for the principal phenotype are asterisked, and SNP-disease associations not previously known are suffixed with a ‘+’. Table S1. shows SNPs associated with T1D (type 1 diabetes).

**Table S3.** SNPs associated with ATD (autoimmune thyroid disease). See legend for Table S2.

**Table S4.** SNPs associated with CEL (celiac disease). See legend for Table S2.

**Table S5.** SNPs associated with MS (multiple sclerosis). See legend for Table S2.

**Table S6.** SNPs associated with NAR (narcolepsy). See legend for Table S2.

**Table S7.** SNPs associated with PBC (primary biliary cirrhosis). See legend for Table S2.

**Table S8.** SNPs associated with PSO (psoriasis). See legend for Table S2.

**Table S9.** SNPs associated with RA (rheumatoid arthritis). See legend for Table S2.

**Table S10.** SNPs associated with UC (ulcerative colitis). See legend for Table S2.

**Table S11.** SNPs associated with CRO (Crohn’s disease). See legend for Table S2.

**Figure S1.** Effect of adjusting 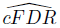 for shared controls. Plots A and B show the ratio between the true 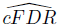 (computed using our method) to the ‘naive’ 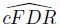 (computed by naively applying the existing split-control approach to shared-control data without adjustment) for a range of p values for the principal phenotype. The p values forming the x-coordinates were obtained from the shared-control design. The left-hand plot shows ratios of true to ‘naive’ cFDR for p values greater than 1 × 10^−10^, demonstrating 2-3 fold underestimation. The right-hand plot shows log-ratios of trueto ‘naive’ cFDR for smaller p values, demonstrating hundred- or thousand-fold underestimation.

**Figure S2.** Summary of pleiotropy between phenotypes. The colour for phenotype *i* (horizontal) and phenotype *j* (vertical) corresponds to the p-value cutoff for significance for phenotype *i*, given that a p-value cutoff for phenotype *j* is less than 5 × 10^−6^.

**Figure S3.** Distribution of log(*M*) amongst null SNPs. *M* is proportional to the variance of the log odds ratio from TDT data, defined as *σ̂*^2^*f* (1 − *f*)), where *f* is the minor allele frequency amongst null SNPs, and *σ̂* is the standard error. Equating the median of *M* with a known expression for variance of the log odds ratio in a case-control study enables back-calculation of the effective number of cases and controls. This technique was used for computing the number of cases and controls in the T1D study, for which p values were obtained from a meta-analysis of case-control and TDT data.

**Figure S4.** Figures S4–S13 are Q-Q plots labeled “*i*|*j*”, where *i* is the principal phenotype and *j* the conditional phenotype. Y axes show 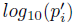; X axes show log quantile (rank) of p values in various sets of SNPs. Each colour corresponds to the Q-Q plot for *p_i_* amongst only SNPs such that *p_j_* is less than a certain cutoff, with the black line corresponding to the Q-Q plot for all SNPs. P values for the principal phenotype are adjusted for the effect of shared controls between studies. A leftward shift with decreasing *p_j_* cutoff indicates enrichment of SNP sets from conditioning on degrees of association with a conditional phenotype, probably due to pleiotropic effects between phenotypes. Because the studies used the ImmunoChip, which covers only potential autoimmune-associated regions, the black line also shows considerable enrichment compared to quantiles. Figure S4 shows Q-Q plots with T1D (type 1 diabetes) as the principal phenotype

**Figure S5.** Q-Q plots with ATD (autoimmune thyroid disease) as the principal phenotype. See legend for Figure S4

**Figure S6.** Q-Q plots with CEL (celiac disease) as the principal phenotype. See legend for Figure S4

**Figure S7.** Q-Q plots with MS (multiple sclerosis) as the principal phenotype. See legend for Figure S4

**Figure S8.** Q-Q plots with NAR (narcolepsy) as the principal phenotype. See legend for Figure S4

**Figure S9.** Q-Q plots with PBC (primary biliary cirrhosis) as the principal phenotype. See legend for Figure S4

**Figure S10.** Q-Q plots with PSO (psoriasis) as the principal phenotype. See legend for Figure S4

**Figure S11.** Q-Q plots with RA (rheumatoid arthritis) as the principal phenotype. See legend for Figure S4

**Figure S12.** Q-Q plots with UC (ulcerative colitis) as the principal phenotype. See legend for Figure S4

**Figure S13.** Q-Q plots with CRO (Crohn’s disease) as the principal phenotype. See legend for Figure S4

